# Proteasome-derived antimicrobial peptides discovered via deep learning

**DOI:** 10.1101/2025.03.17.643752

**Authors:** Xiaoqiong Xia, Marcelo D. T. Torres, Cesar de la Fuente-Nunez

## Abstract

Recent computational discoveries have identified numerous bioactive peptides within the human proteome, as well as across the broader tree of life, which were previously unrecognized for their roles in host immunity. These findings have led us to propose the “cross-talk hypothesis”, suggesting that many molecules, such as proteins and peptides, traditionally viewed as extraneous to immune function may in fact actively contribute to immunity. Building on our earlier studies, which uncovered proteasome-derived peptides with putative antimicrobial activity in the human proteome, here we systematically interrogated the proteasome, a large protein complex responsible for degrading and recycling damaged or surplus proteins, for additional antimicrobial peptides. Using deep learning, we systematically mined ProteasomeDB, a curated repository of proteasomal cleavage and splicing events, to predict antibiotic activity against 11 clinically relevant pathogens. This deep learning approach uncovered 59 candidate peptides (“proteasomins”) with a median minimum inhibitory concentration (MIC) of ≤64LμmolLLL^1^. Refinement yielded 21 sequence-diverse proteasomins, which were characterized for their physicochemical properties. These peptides were enriched in cationic residues and exhibit enhanced amphiphilicity, key attributes for disrupting microbial membranes. Dimensionality reduction via UMAP further showed that proteasomins are sequence-distinct from known antimicrobial peptides, underscoring their novelty and potential for unique mechanisms of action. Moreover, comparative analyses revealed that cis- and trans-spliced proteasomins exhibit similar predicted antimicrobial activities, suggesting that critical structural determinants remain conserved irrespective of splicing modality. Collectively, these findings expand our understanding of the proteasome, underscore the extensive, previously unrecognized repertoire of innate immune peptides, and provide a promising foundation for developing innovative therapeutics to combat multidrug-resistant pathogens.

## Introduction

Recent computational approaches have begun to illuminate a vast and previously unrecognized landscape of bioactive peptides hidden within proteins across diverse organisms^1-7^. Indeed, we recently proposed the “cross-talk hypothesis,”^1^ to illustrate how certain molecules—once dismissed as extraneous to immune function—may in fact contribute to host immunity. Evidence suggests that these peptides can arise via proteolytic cleavage events from larger precursor proteins^5,7^, subsequently playing critical roles in host defense^8^.

Our computational exploration of the human proteome^1,7^ revealed thousands of novel peptides with antibiotic activity, several of which originated from the proteasome. Building on these earlier findings, we hypothesized that the proteasome would be an excellent source of antimicrobial petpides, despite the historical assumption that it only produced “junk” fragments destined for antigen presentation or degradation. The proteasome, an intracellular protein degradation complex, is best known for recycling damaged or surplus proteins and generating peptide fragments for presentation via major histocompatibility complex class I (MHC I) molecules^9^.

Given converging evidence pointing to the proteasome’s broader immunological relevance, proteasomins represent a promising yet underexplored reservoir of antimicrobial molecules. We hypothesized that proteasomes might undergo functional or compositional shifts upon microbial infection, increasing the production of antimicrobial peptides. By applying a deep learning approach to a large corpus of proteasomal peptides, we have identified a suite of novel antimicrobial peptides—”proteasomins”—that expand our understanding of the proteasome’s role in immunity and reinforce the cross-talk hypothesis.

## Results

### Mining antibiotic candidates from proteasome-derived sequences

To identify candidate antimicrobials systematically, we established an integrated computational screening pipeline (**Figure 1**). First, cis- and trans-spliced peptides were extracted from ProteasomeDB^10^, a curated database of proteasomal cleavage and splicing events. To enhance both the likelihood of biological activity and synthetic feasibility, we restricted our search to peptides shorter than 50 amino acid residues, resulting in a final set of 15,784 unique sequences^11^. These sequences were subsequently evaluated using our APEX deep-learning model^12^, which predicts minimum inhibitory concentration (MIC) values against 11 clinically significant bacterial pathogens. The median MIC cutoff was set as 64 *μ*mol L^-1^.

**Figure 1.**
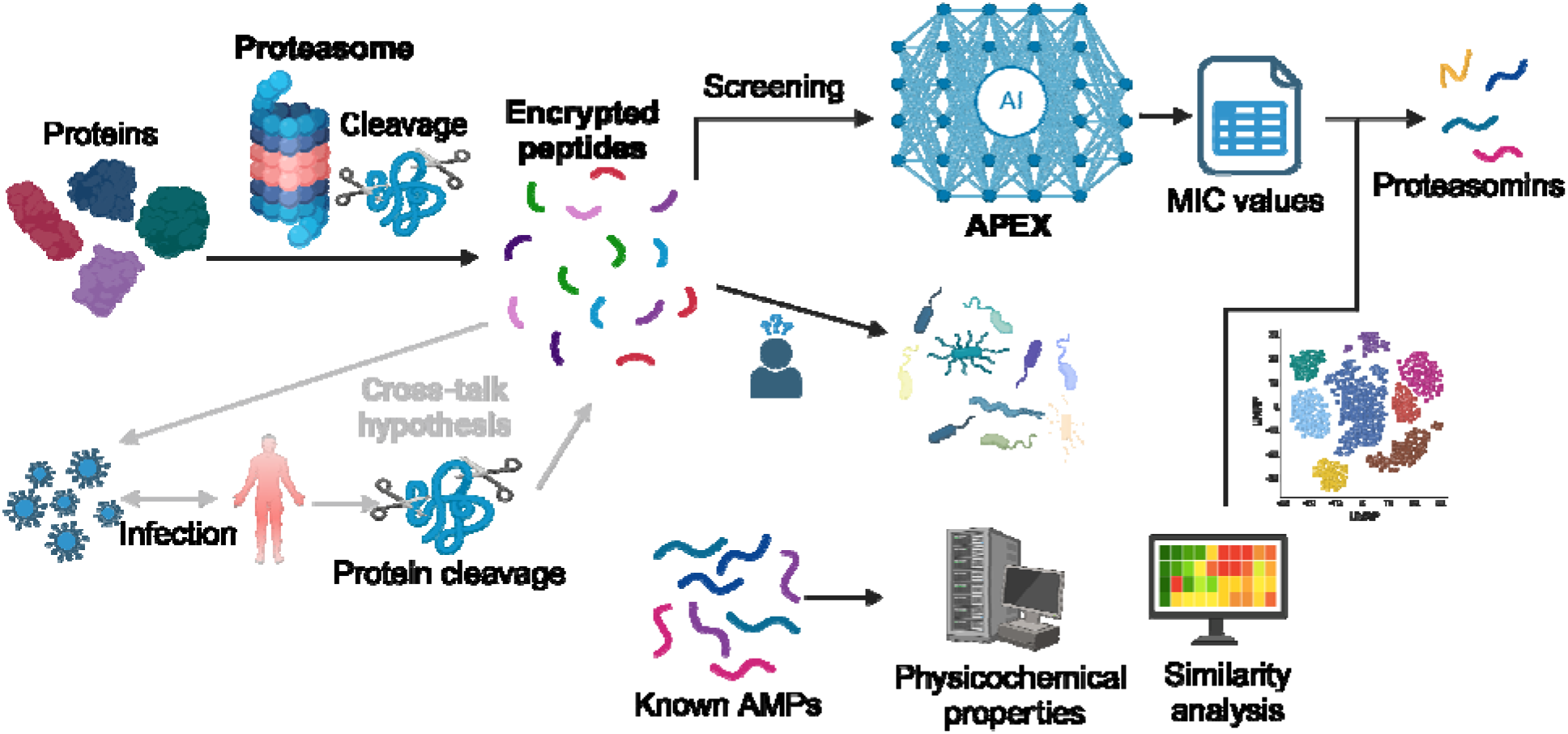
Computational framework for the discovery of antimicrobial peptides in the proteasome. Host proteins are proteolytically processed by the proteasome under both normal and infection conditions, generating “encrypted peptides” that may exhibit antimicrobial activity (the “cross-talk hypothesis”). The APEX deep-learning model screens these peptides for their predicted minimal inhibitory concentration (MIC) against clinically relevant pathogens. Peptides meeting the MIC cutoff (≤64□μmol□L□^1^) are designated as “proteasomins.” Comparative analyses—encompassing known antimicrobial peptides (AMPs), physicochemical profiling, and dimensionality reduction (UMAP)—further refine and characterize proteasomins, highlighting their distinct sequence space and potential as novel therapeutic agents.

### Identification of proteasomins and their antimicrobial potency

APEX evaluated the 15,784 peptide sequences for antimicrobial activity across 11 clinical relevant pathogens, including the species from the World Health Organization (WHO) watchlist, the ESKAPEE (*Enterococcus faecium, Staphylococcus aureus, Klebsiella pneumoniae, Acinetobacter baumannii, Pseudomonas aeruginosa, Escherichia coli*, and *Enterobacter spp)*. Candidate peptides were ranked based on median MIC values. Using the cutoff of median MIC ≤64 *μ*mol L^-1^, we obtained 59 peptides (**Table S1**), emphasizing the antimicrobial potential of PDEPs.

### Composition of proteasomins

The analysis of the amino acid composition of the proteasome-derived antimicrobial peptides (proteasomins) identified by APEX revealed distinct patterns compared to peptides listed in the ProteasomeDB and traditional AMPs from databases (**Figure 2a**). Amino acid residues such as lysine (K), arginine (R), and glycine (G), traditionally associated with antimicrobial activity due to their cationic and flexibility, were enriched in proteasomins relative to the general proteasomal peptide pool in ProteasomeDB. Interestingly, proteasomins exhibited a notable increase in the frequency of cationic residues (K and R) compared to the broader dataset, indicating potential for enhanced electrostatic interactions with negatively charged bacterial membranes, a marked characteristic of potent antimicrobial peptides. Conversely, the acidic residues, glutamic acid (E) and aspartic acid (D), typically less favorable for antimicrobial activity, showed lower frequencies within the proteasomin group compared to ProteasomeDB, aligning more closely with known AMPs and probably why they were ranked as antimicrobials by APEX.

**Figure 2.**
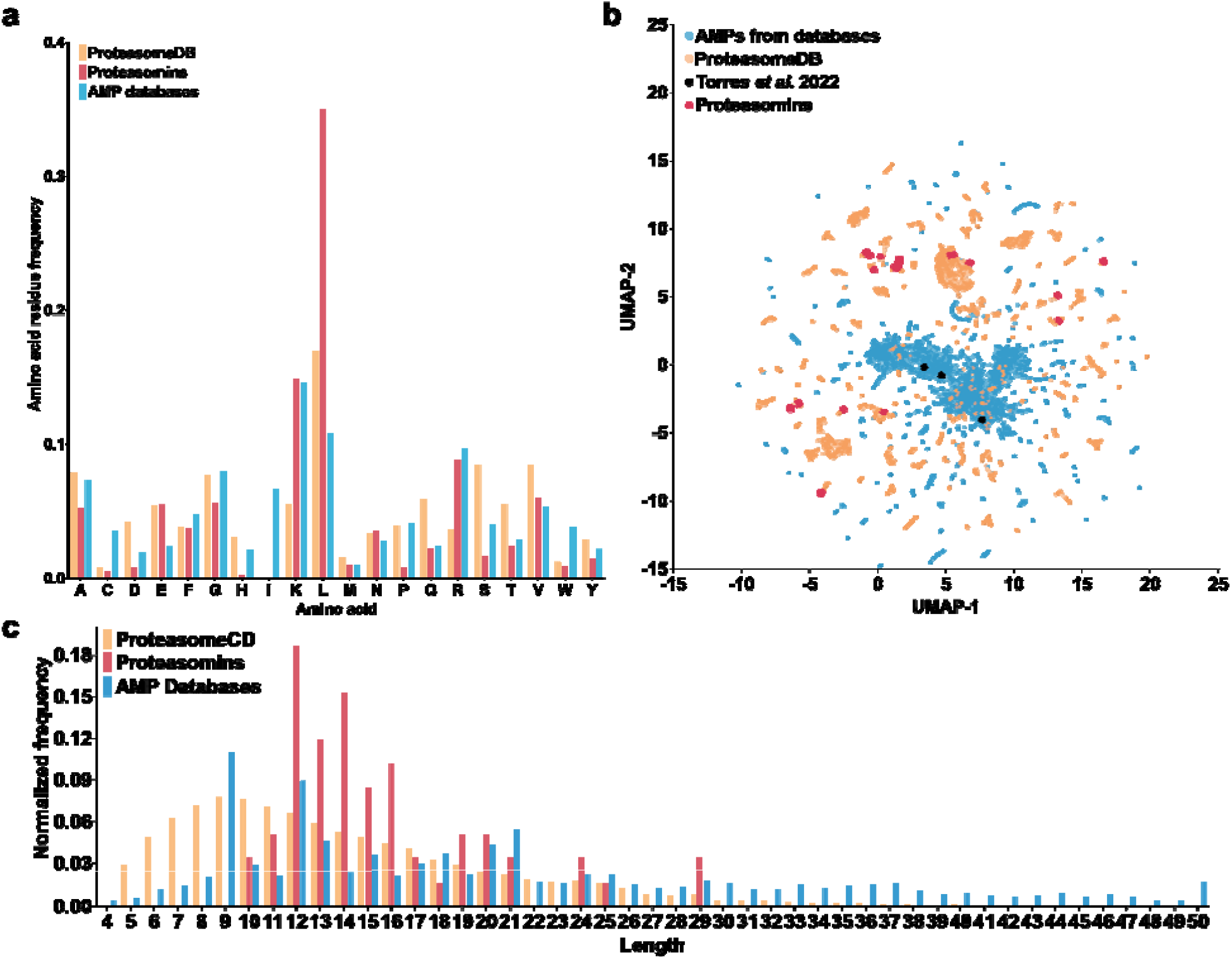
Compositional and sequence-space characteristics of proteasomins. **(a)** Amino acid residue frequencies of proteasomins, compared against peptides in ProteasomeDB and known AMPs databases (DBAASP, APD3, and DRAMP 3.0). Proteasomins exhibit notable enrichment of cationic residues (e.g., K, R) and aliphatic residue (L), aligning with features often associated with antimicrobial activity. However, glutamic acid (E) is also overrepresented as in the ProteasomeDB sequences. **(b)** UMAP-based visualization of peptide sequence space. Points represent peptides from AMP databases (blue), ProteasomeDB (orange), previously characterized proteasome-derived AMPs (black, from Torres et al. 2022), and APEX-discovered proteasomins (red). Each peptide is represented as a vector of sequence similarity scores that capture its relationship with every other peptide in the dataset. These high-dimensional vectors are then projected into a two-dimensional space using UMAP, revealing that proteasomins occupy a distinct region of sequence space compared to known AMPs. **(c)** Normalized length distributions for proteasomins (red) in comparison to ProteasomeDB (orange) and AMP databases (blue). Proteasomins identified by APEX predominantly range from 8 to 20 residues, a length conducive to membrane disruption and antimicrobial efficacy.

A particularly striking observation was the complete absence of isoleucine (I) residues in both ProteasomeDB peptides and, consequently, in the proteasomins. Isoleucine is a branched-chain hydrophobic amino acid commonly found in traditional AMPs, where it contributes to hydrophobic interactions crucial for membrane insertion and disruption. However, excessive hydrophobicity, especially from residues like isoleucine, can promote peptide aggregation, decrease solubility, and increase the potential for non-specific interactions with host membranes, leading to cytotoxicity. The absence of I residues in proteasomins may reflect an evolutionary and functional constraint that favors maintaining solubility and reducing aggregation potential, thereby optimizing these peptides for selective antimicrobial activity. This absence may also be attributed to the proteasome’s cleavage preferences, which might avoid generating fragments with isoleucine at key positions, possibly as a strategy to prevent the formation of aggregation-prone sequences.

### Sequence similarity of proteasomins compared to known AMPs

Following the application of a similarity filter (cutoff = 0.7; see **Peptide ranking and filtering** in **Methods**), we selected 21 proteasomins that exhibited low similarity to both known AMPs and other proteasomin sequences (**Table S2**). To visualize the sequence relationships among our newly identified proteasomins, previously characterized antimicrobial peptides (AMPs), AMPs identified as proteasome-derived AMPs, and a the whole ProteasomeDB dataset, we performed a uniform manifold approximation and project (UMAP) analysis (**Figure 2b**). The proteasomin sequences (red points) are widely dispersed, indicating high sequence diversity and minimal clustering among themselves. They do not overlap with the cluster of known AMPs (blue points), suggesting that proteasomins are sequencely distinct from established AMP families. In contrast, the proteasomins predicted by a physicochemical features-based scoring function^7^ (black points; **Table S3**) show compelte overlap with known AMPs, as expected since the premises for their discovery were few properties that are marked features of AMPs. Curiously, those three sequences are considerably different from the APEX-discovered proteasomins. This distribution pattern underscores the novelty of our proteasomin sequences and highlights their low sequence similarity to known AMPs. The distinct clustering of proteasomins implies the discovery of a new class of proteasome-derived peptides with potentially unique antimicrobial features due to their different composition. Such diversity expands our understanding of the proteasome-derived peptides as a source of novel therapeutic candidates and provides a strong foundation for future functional studies aimed at harnessing these peptides to address emerging infectious disease challenges.

Further characterization of peptide length distributions showed that proteasomins predominantly ranged from 8 to 20 amino acids, a length similar to peptides frequently present in AMP databases and optimal for membrane permeability and microbial targeting (**Figure 2c**). Compared to ProteasomeDB, which included peptides spanning a broader length distribution up to 50 amino acid residues, proteasomins presented a narrower and distinctly peaked length distribution profile centered around shorter, bioactive lengths (12-16 residues long). Such distribution implies a selection pressure or functional constraint towards lengths conducive to antimicrobial activity created by APEX.

Collectively, these observations highlight the unique biochemical signatures of proteasomins and underscore their potential physiological roles in innate immunity. By contrasting amino acid composition and peptide length characteristics between proteasomins, ProteasomeDB, and known AMPs, our findings reinforce the hypothesis that proteasome-derived peptides represent an evolutionary conserved and underexplored source of natural antimicrobial agents.

### Physicochemical properties of proteasomins

To delve deeper into the properties of proteasomins MPs, we analyzed their physicochemical properties using the DBAASP tool^13^. Properties such as net charge (**Figure 3a**), normalized hydrophobicity (**Figure 3b**) and hydrophobic moment (**Figure 3c**), amphiphilic index (**Figure 3d**), isoelectric point (**Figure 3e**), linear moment (**Figure 3f**), and aggregation (**Figure 3g**) and disoredered confirmation (**Figure 3h**) propensities were calculated for known AMPs, the whole ProteasomeDB, and the newly predicted proteasomins.

**Figure 3.**
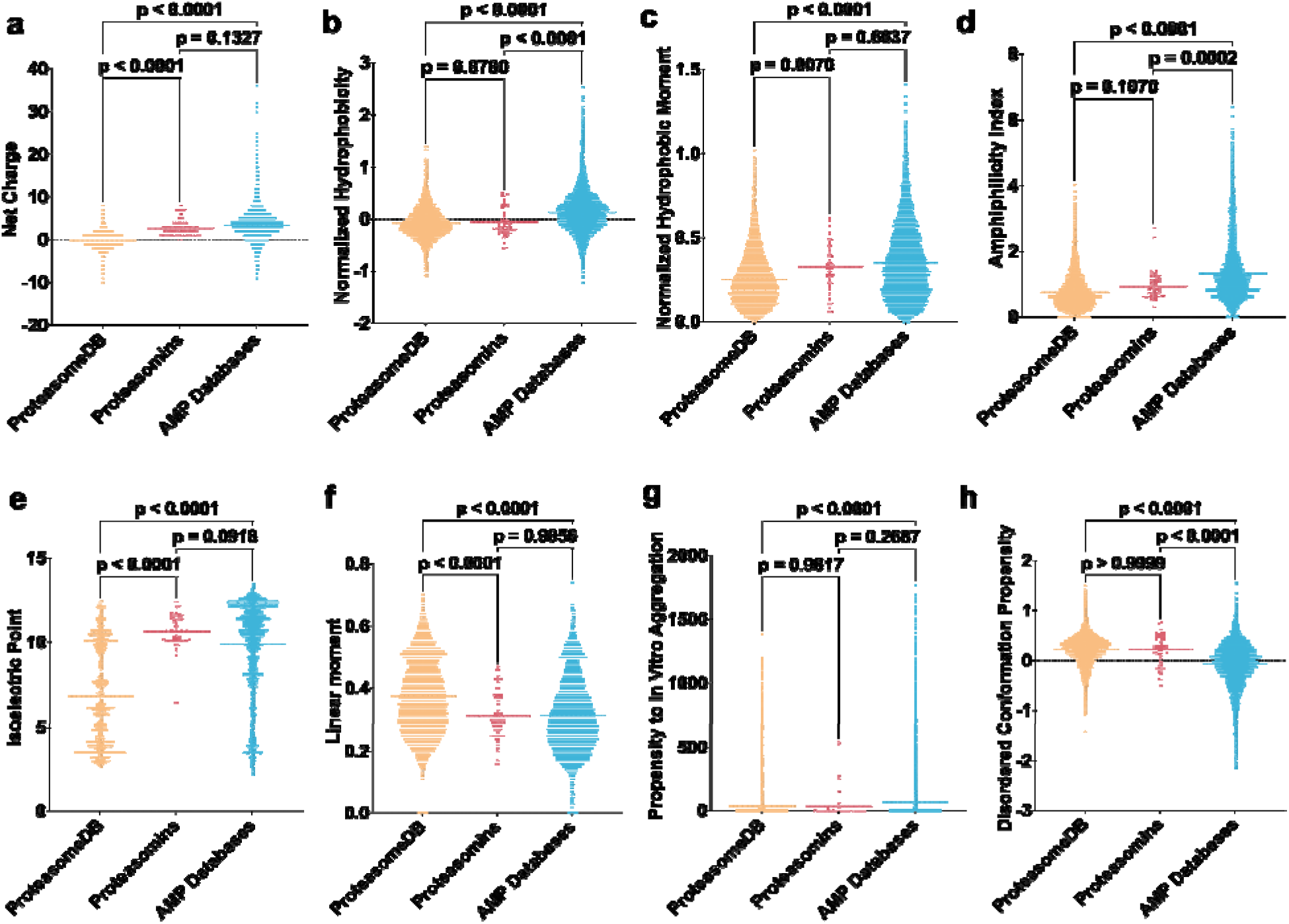
Physicochemical properties of proteasomins compared with ProteasomeDB peptides and known AMPs. Each violin plot shows the distribution of a key physicochemical characteristic across three datasets: ProteasomeDB (orange), proteasomins identified by APEX (red), and AMPs from databases (blue). **(a)** Net charge, critical for electrostatic interactions with negatively charged bacterial membranes. **(b)** Normalized hydrophobicity, indicating the balance of polar and nonpolar residues that governs membrane insertion. **(c)** Normalized hydrophobic moment, **(d)** amphiphilicity index, both closely tied to membrane-disruptive activity and mechanism of action. **(e)** Isoelectric point, influencing how peptides behave in various pH environments and interact with microbial surfaces. **(f)** Linear moment, a measure of the spatial arrangement of hydrophobic/polar segments that can affect membrane targeting. **(g)** Propensity to aggregate *in vitro*, which has implications for peptide stability and toxicity. **(h)** Disordered conformation propensity, reflecting a peptide’s flexibility and potential for selective targeting. Statistical significance was assessed using two-tailed t-tests followed by the MannWhitney test; p values are shown in the graph. The solid line within each violin plot represents the mean value obtained for each group.

Proteasomins displayed distinctive physicochemical properties relative to peptides cataloged in ProteasomeDB and AMP databases, suggesting their optimized potential for antimicrobial activity. Our analysis revealed that proteasomins have significantly elevated amphiphilicity compared to ProteasomeDB peptides (p < 0.0001), aligning closely with traditional antimicrobial peptides, which often utilize amphiphilic helices for membrane disruption. Furthermore, proteasomins demonstrated significantly higher normalized hydrophobic moments than those in ProteasomeDB (p < 0.0001), supporting their capability to interact effectively with bacterial membranes.

A crucial determinant of peptide function, the net charge of proteasomins was significantly higher relative to peptides listed in ProteasomeDB (p < 0.0001) but did not differ significantly from peptides in AMP databases (p = 0.1327), reinforcing their predicted antimicrobial function through electrostatic interactions with negatively charged bacterial membranes. Moreover, proteasomins exhibit higher isoelectric points relative to ProteasomeDB (p < 0.0001), mirroring the strongly basic nature characteristic of known antimicrobial peptides and favoring interactions in microbial environments and also a reason for the selection by APEX.

In contrast, other physicochemical attributes, including normalized hydrophobicity and linear moments, were comparable between proteasomins and ProteasomeDB peptides, suggesting intrinsic hydrophobic features consistent across proteasomal products. Additionally, the propensity for adopting disordered conformations and *in vitro* aggregation did not significantly differ between proteasomins and ProteasomeDB peptides, although proteasomins significantly deviated from AMP databases (p < 0.0001). This indicates that despite similarities in hydrophobic properties and aggregation tendencies with other proteasomal peptides, proteasomins possess specialized physicochemical properties that could selectively enhance antimicrobial functionality without adverse aggregative side effects.

These findings collectively emphasize that proteasomins possess physicochemical properties uniquely suited for antimicrobial activity, distinguishing them from general proteasomal peptides and positioning them as valuable components within innate immunity.

We then employed UMAP for dimensionality reduction^14^ (**Figure 4**). Proteasomins share certain physicochemical properties with broader ProteasomeDB peptides and known AMPs. This strong similarity suggests that these proteasomins may share key functional traits, such as charge distribution, hydrophobicity, and structural motifs, which are often associated with antimicrobial activity. Consequently, the computational evidence supports the hypothesis that these candidates could exhibit antimicrobial properties. The proteasomins also lie within or near the main ProteasomeDB cluster, these peptides retain properties characteristic of the general set of ProteasomeDB sequences, thus they are not outliers. Proteasomins still belong to the broader PDEP family, in terms of overall composition and sequence features.

**Figure 4.**
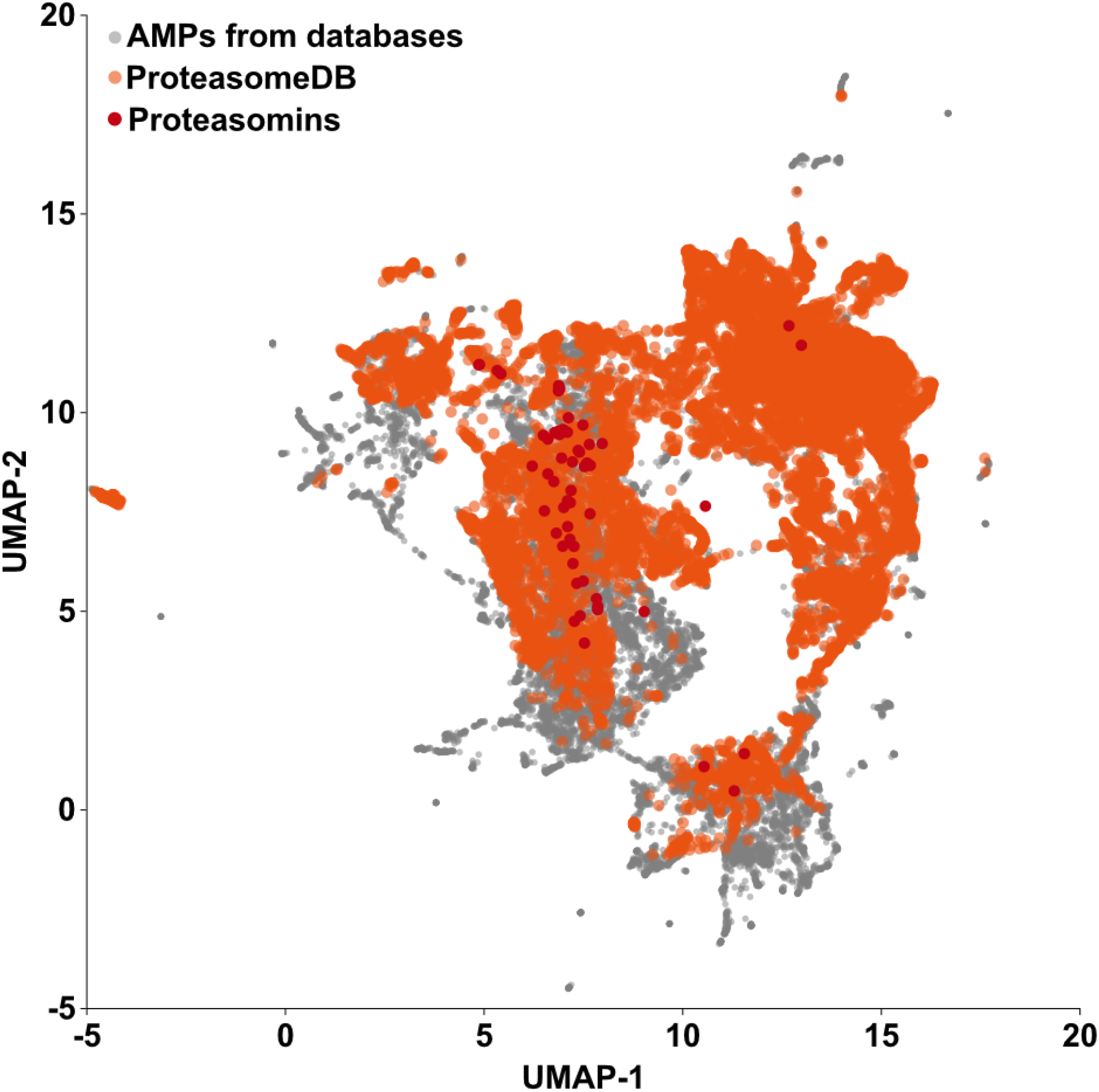
Two dimensional visualization of proteasomins within physicochemical feature space. The scatterplot shows peptide sequences from AMP databases (DBAASP, APD3, DRAMP 3.0; gray), ProteasomeDB (orange), and the 59 proteasomins identified by APEX (red). Each peptide’s physicochemical properties (e.g., charge, hydrophobicity, amphiphilicity) were reduced to two dimensions using UMAP. This approach demonstrates that the newly discovered proteasomins exhibit physicochemical properties similar to those of both known AMPs and general proteasome-derived peptides.

### Comparative Analysis of Antimicrobial Activity Between Cis- and Trans-Spliced Peptides

Among the 59 identified proteasomins, 12 were trans-spliced and 10 were cis-spliced by proteasomes, the remaining were undefined. Comparison of the predicted median MIC values between cis- and trans-spliced proteasomins revealed no significant difference (Wilcoxon rank-sum test, p = 0.62), indicating that both splicing mechanisms produce peptides with similarly high antimicrobial activity. Analysis of the broader proteasomeDB dataset, which includes 4,064 trans-spliced and 2,869 cis-spliced peptides, showed a significant difference in the overall MIC distribution (Kolmogorov–Smirnov test, p = 0.0001); however, the median MIC values between the two groups were statistically comparable (Wilcoxon rank-sum test, p = 0.22). These findings suggest that, despite variability in the MIC distribution, the central antimicrobial efficacy is preserved regardless of the splicing mechanism. This conservation implies that the structural and functional determinants critical for antimicrobial activity are maintained in both cis- and trans-spliced peptides, underscoring the potential of each splicing pathway in generating effective antimicrobial agents.

## Discussion

In this work, we used an integrated computational approach to uncover proteasome-derived antimicrobial peptides, expanding the cross-talk hypothesis to encompass an underexplored reservoir of innate immune factors. Starting with 15,784 peptides from ProteasomeDB, we restricted our search to sequences under 50 residues. The APEX deep-learning model identified 59 “proteasomins” with median MIC values ≤64LμmolLL^-1^, 21 of which passed a stringent sequence similarity filter to ensure novelty.

Proteasomins display distinct biochemical signatures compared to both the broader ProteasomeDB dataset and AMP databases. They are enriched in cationic and amphiphilic residues, a hallmark of membrane-acting antimicrobials. The absence of isoleucine (I) suggests that these peptides have evolved—or have been selectively generated—to minimize aggregation and toxicity risks. Dimensionality reduction via UMAP emphasizes their uniqueness, showing that proteasomins diverge from known AMP clusters.

Furthermore, proteasomins exhibit comparable predicted antimicrobial activity regardless of whether they originate from cis- or trans-splicing events. This finding indicates that splicing modality does not undermine their ability to target pathogens. Overall, our results point to the proteasome not just as a generator of antigenic peptides but also as a potential source of functional molecules involved in innate defense.

The discovery of proteasomins underscores the possibility that many proteasomal byproducts, previously viewed as “junk,” may play unappreciated roles in host immunity. Experimental validation of proteasomin activity—along with *in vivo* efficacy and mechanism-of-action studies—will be crucial to harnessing their therapeutic potential in the face of mounting antimicrobial resistance.

## Conclusion

In summary, our integrated computational pipeline uncovered a novel class of proteasome-derived antimicrobial peptides, termed proteasomins, which show predicted activity against 11 clinically relevant bacterial pathogens. By applying the APEX deep-learning model to ProteasomeDB, we identified 59 such peptides (median MIC ≤64LμmolLL^-1^), refining these down to 21 sequence-diverse candidates. Notably, proteasomins feature a distinctive amino acid composition (e.g., enriched cationic residues, absence of isoleucine) and appear in a distinct region of sequence space relative to known AMPs and other proteasome-derived fragments. Cis- or trans-splicing did not significantly influence their antimicrobial potency.

Collectively, these findings support the cross-talk hypothesis by demonstrating that the proteasome—beyond its canonical functions in antigen processing—serves as an underexplored reservoir of encrypted peptides that may directly contribute to innate immunity. Proteasomins thus expand the repertoire of natural antimicrobial agents and present compelling avenues for next-generation anti-infective strategies. Future experimental and translational studies are warranted to fully exploit their clinical potential and deepen our understanding of proteasome-mediated host defense.

## Methods

### Data Collection from ProteasomeDB

We retrieved cis- and trans-spliced peptides from ProteasomeDB, a proteomic database documenting validated proteasomal cleavage and peptide splicing events^10^. Peptides ranging from 5 to 66 amino acids were initially collected, after which those longer than 50 amino acid resoidues were excluded to facilitate synthesis and functional characterization. The final data corpus included 15,784 unique sequences, comprising both spliced and nonspliced peptides.

### AMP Prediction Using APEX

All candidate peptides were screened for antimicrobial activity using our APEX deep-learning model^2^. APEX integrates recurrent neural network architectures trained on curated public AMP databases (e.g., DBAASP^13^) and in-house MIC datasets. The model predicted MIC values against 11 clinically relevant pathogens, including *E. coli* ATCC 11775, *P. aeruginosa* PAO1, *P. aeruginosa* PA14, *S. aureus* ATCC 12600, *E. coli* AIC221, *E. coli* AIC222, *K. pneumoniae* ATCC 13883, *A. baumannii* ATCC 19606, methicillin-resistant *S. aureus* ATCC BAA-1556, vancomycin-resistant *E. faecalis* ATCC 700802 and vancomycin-resistant *E. faecium* ATCC 700221, after which peptides were ranked by median MIC <64 *μ*mol L^-1^. Peptides were ranked based on their median MIC.

### Similarity Analysis with Known AMPs and UMAP Visualization

To assess the novelty of these PDEPs, we compared their sequence similarity with a comprehensive database of 19,762 known AMPs from DBAASP^13^, APD3^15^, and DRAMP 3.0^16^ public databases. The similarity calculation employed pairwise sequence alignment using the BLOSUM50 substitution matrix implemented in the scikit-bio library (skbio.alignment._pairwise). Each PDEPs was compared individually against all known AMP sequences to determine the maximum similarity score. Finally, a similarity matrix was constructed to quantify the pairwise sequence similarities among all the known AMPs and proteasome-drived AMPs. Each peptides are characterized with the similarity vector. To visualize the relationships and clustering patterns among these peptides, dimensionality reduction was performed using the Uniform Manifold Approximation and Projection (UMAP) technique. The resulting two-dimensional embeddings (UMAP-1 and UMAP-2) provided a visual representation of peptide similarities and facilitated the identification of distinct peptide clusters based on sequence characteristics.

### Peptide Ranking and Filtering

The identified PDEPs were ranked based on predicted MIC in decrease order (highest to lowest activity potential). Subsequently, peptides exhibiting greater than 70% sequence similarity with any known AMP were excluded to ensure the novelty of the candidate peptides. Additionally, peptides demonstrating internal redundancy, defined as having greater than 70% similarity to any other peptide within the proteasome-derived AMP set, were also removed. This rigorous filtering approach ensured the selection of unique and novel candidate peptides with low similarity both to known AMPs and within the dataset.

### Physicochemical Property Analysis Using DBAASP Tool

The DBAASP tool^13^ was used to calculate 12 physicochemical descriptors, including normalized hydrophobic moment, normalized hydrophobicity, net charge, isoelectric point, penetration depth, tilt angle, disordered conformation propensity, linear moment, propensity to *in vitro* aggregation, angle subtended by the hydrophobic residues, amphiphilicity index, propensity to PPII coil for each peptide. These values formed a feature set used for subsequent clustering and visualization.

### Statistical Analysis

Predicted MIC values for cis- and trans-spliced peptide groups were compared using the Wilcoxon rank-sum test, and differences in their overall MIC distributions were evaluated via the Kolmogorov–Smirnov (K–S) test, with significance set at p <0.05. MIC values are presented as medians with interquartile ranges. All statistical analyses were performed using Python.

## Acknowledgements

Cesar de la Fuente-Nunez holds a Presidential Professorship at the University of Pennsylvania and acknowledges funding from the Procter & Gamble Company, United Therapeutics, a BBRF Young Investigator Grant, the Nemirovsky Prize, Penn Health-Tech Accelerator Award, Defense Threat Reduction Agency grants HDTRA11810041 and HDTRA1-23-1-0001, and the Dean’s Innovation Fund from the Perelman School of Medicine at the University of Pennsylvania. Research reported in this publication was supported by the Langer Prize (AIChE Foundation), NIH R35GM138201, and DTRA HDTRA1-21-1-0014.

